# Repurposing the atypical Type I-G CRISPR system for bacterial genome engineering

**DOI:** 10.1101/2023.04.24.538059

**Authors:** Qilin Shangguan, Malcolm F White

## Abstract

The CRISPR-Cas system functions as a prokaryotic immune system and is highly diverse, with six major types and numerous sub-types. The most abundant are type I CRISPR systems, which utilise a multi-subunit effector, Cascade, and a CRISPR RNA (crRNA) to detect invading DNA species. Detection leads to DNA loading of the Cas3 helicase-nuclease, leading to long range deletions in the targeted DNA, thus providing immunity against mobile genetic elements (MGE). Here, we focus on the type I-G system, a streamlined, 4- subunit complex with an atypical Cas3 enzyme. We demonstrate that Cas3 helicase activity is not essential for immunity against MGE *in vivo* and explore applications of the *Thioalkalivibrio sulfidiphilus* Cascade effector for genome engineering in *Escherichia coli*. Long range, bidirectional deletions were observed when the *lacZ* gene was targeted. Deactivation of the Cas3 helicase activity dramatically altered the types of deletions observed, with small deletions flanked by direct repeats that are suggestive of microhomology mediated end joining. When donor DNA templates were present, both the wild-type and helicase deficient systems promoted homology-directed repair (HDR), with the latter system providing improvements in editing efficiency, suggesting that a single nick in the target site may promote HDR in *E. coli* using the type I-G system. These findings open the way for further application of the type I-G CRISPR systems in genome engineering.

## Introduction

CRISPR systems (classified into types I-VI) function in adaptive immune defence in prokaryotes (1, 2). The programmable, sequence specific nature of CRISPR effectors has led to widespread repurposing of CRISPR for genome editing (3). In particular, the single subunit effectors Cas9 (type II) (4, 5) and Cas12a (type V) have been intensively studied and applied in genome editing over the last decade. Type I CRISPR systems are more complex, with multi-subunit effectors, and have been less widely applied in gene editing applications, despite that fact that they are the most prevalent system in bacteria (7).

The signature enzyme of type I systems is Cas3, a helicase-nuclease fusion that degrades double strand DNA once recruited to specific sites by the Cascade complex (7, 8). The broad distribution of type I CRISPR systems in bacteria and archaea has potentiated prokaryotic genome editing applications using the species’ own endogenous CRISPR apparatus. Examples include the type I-A system in *Saccharolobus islandicus (9)*, type I-B in *Clostridia* (10, 11, 12) and *Haloarcula hispanica* (13), type I-C in *Pseudomonas aeruginosa* (14), types I-E (15, 16, 17, 18, 19) and I-F (20, 21, 22, 23) in a range of species and type I-G in *Bifidobacteria* (24). Examples of genome engineering in heterologous hosts are less common, but types I-C and I-F from *Pseudomonas* have been used in this manner (14, 25). There are also successful attempts utilizing type I CRISPR in eukaryotic systems (26, 27, 28, 29, 30, 31, 32, 33), showing promising potential for gene deletion and modulation of gene expression.

These genome engineering approaches typically result in long-range genomic deletions, due to the processive nature of Cas3. Cas3 is a dual helicase and nuclease protein that plays a key role in target DNA degradation (34, 35, 36). It is either recruited by the Cascade complex upon binding target dsDNA (35, 37, 38), or pre-associated with Cascade and allosterically activated upon dsDNA recognition (26, 39). Cas3 is a superfamily 2 helicase that unwinds target DNA in a 3’ to 5’ direction and degrades the resultant ssDNA in the HD nuclease site, leading to a 3’ to 5’ non-target strand degradation (36, 40, 41, 42). Strand switching, or loading of a second Cas3 on the target strand, can lead to bi-directional DNA degradation (38, 43).

Previously, we reported the structure and mechanism of the type I-G system from *Thioalkalivibrio sulfidiphilus* (39), demonstrating the constitutive association of Cascade with the Cas3 subunit. Here, we repurpose the *T. sulfidiphilus* system for gene disruption in *E. coli* via long-range, bidirectional DNA degradation, and show that abolition of the helicase activity of *T. sulfidiphilus* Cas3 results in efficient gene disruption by generation of small deletions in target genes.

## Methods

### Cloning

For *E. coli* genome editing, the pM2 vector was constructed based on the previously described pACE-M1 (39). The original T7 promoter was replaced by an araBAD promoter using overlap PCR extension (39) with overlap primers. Restriction sites (*NcoI* and *SalI*) were introduced to facilitate further construction. The *cas3* gene was digested with *Nco*I and *Sal*I (Thermo Scientific) and ligated into the promoter-swapped vector to generate the pM2 vector. Site directed mutagenesis of the *cas3* gene in pM2 was carried out using standard protocols with Phusion enzyme (Thermo Scientific). Two *Bpi*I restriction sites with type I-G repeat sequence were introduced to pRAT-Duet MCS-I to get the spacer replaceable backbone of the pSPACER plasmid. 5’-phosphorylated oligos of CRISPR spacers were annealed and ligated into the *Bpi*I digested pSPACER backbone to obtain the pSPACER vector with target spacer (39).

For experiments that required a DNA donor, we constructed the pHR vector by introducing homologous template into the pSPACER vector. Two 615 bp homologous arms (donor) for homologous directed repair (HDR) were PCR amplified from the *E. coli* MG1655 genome. The donor was incorporated into pSPACER MCS-II using restriction sites *Nde*I and *Avr*II and ligation. All final constructs were verified by sequencing (GATC Biotech, Eurofins Genomics, DE). Primers and synthetic genes were obtained from Integrated DNA Technologies (Coralville, IA, USA), sequence can be found in **Supplementary table 1 and table 2**.

### Genome targeting by the type I-G CRISPR system

pM2 was transformed into *E. coli* MG1655. Transformants were selected using 100 µg ml^−1^ ampicillin. Competent cells were prepared by diluting an overnight culture 50-fold into fresh, selective LB medium. The culture was incubated at 37 °C, 220 rpm to reach OD_600_ 0.4 to 0.5. Cells were collected by centrifugation and the pellet resuspended in an equal volume of 100 mM CaCl_2_, 40 mM MgSO_4_. Following incubation on ice for 30 min, cells were collected and resuspended in 0.1 volumes of the same buffer containing 10 % glycerol. Aliquots were stored at -80 °C. 60 ng pSPACER or pHR was transformed into 60 µl competent cells. 400 µl LB medium was added after heat shock and cells incubated at 37 °C for 80 min. 100 µl aliquots of cells were applied onto 10 cm petri dishes in a 10-fold serial dilution for colony number counting and the number was corrected for dilution and volume to obtain colony-forming units (cfu) 0.1ml^−1^. The LB agar plates contained 100 µg ml^−1^ ampicillin, 12.5 µg ml^−1^ tetracycline, 1 mM IPTG, 0.2 mg/ml X-gal and 0.2 % (w/v) L-arabinose for induced plates. Further details are available in **Supplementary table 3**.

### Tiling PCR

Transformants were submitted for colony PCR with sets of primers (**Supplementary table 1**). 10 µl MyTaq™ Red Mix (Bioline, Meridian bioscience) was used with 2 µl 20 µM primer mix, colonies were added into the reaction. PCR products were analysed by separation on a 0.8 % agarose gel, running in 1 X TBE buffer.

### Plasmid challenge assay

The method was described previously (39). Briefly, pACE-M1 (Amp^R^) encompassing the *cas7, cas8g* and *csb2* genes was co-transformed into *E. coli* C43 (DE3) strain with pCDF (Spc^R^) vector containing a CRISPR array that targets tetracycline resistance gene (Tet^R^). *cas3* or *cas3* mutant gene in pRAT-Duet vector with Tet^R^ was used to activate type I-G interference. Transformation reactions were then applied to three different selective plates in a 10-fold dilution series to investigate type I-G targeting. LB agar plates were supplemented with 100 µg ml^−1^ ampicillin and 50 µg ml^−1^ spectinomycin when selecting for recipients only; transformants were selected on LB agar containing 100 µg ml^−1^ ampicillin, 50 µg ml^−1^ spectinomycin, 25 µg ml^−1^ tetracycline, and further supplemented with 0.2 % (w/v) D-lactose and 0.2 % (w/v) L-arabinose for full induction of the type I-G system. Plates were incubated at 37 °C for 16–18 h. statistical analysis was performed with Prism8 (GraphPad). The experiment was performed with two biological replicates and at least two technical replicates.

### Phage immunity assay

The method was described previously (39). Briefly, the type I-G system was built in *E. coli* C43(DE3) by introducing pACE-M1, pCDF-*lpa* (CRISPR array targeting the phage P1 *lpa* gene) and pRAT-Cas3 plasmids. Cells were cultured overnight and infected with phage P1 at a range of MOI (multiplicity of infection) in 96-well plates. The OD_600_ of the culture in the plate was measured using a FilterMax F5 Multi-Mode Microplate Reader (Molecular Devices) every 20 min over 20 h. The experiment was carried out with two biological replicates and three technical replicates. The OD_600_ was plotted against time using Graphpad Prism 8.

## Results

### Structure analysis of type I-G Cas3

The type I-G CRISPR system from *Thioalkalivibrio sulfidiphilus* consists of four protein-coding genes and a CRISPR locus. We previously reported a biochemical and structural study of this complex (39), demonstrating that Cas3 is an integral component of this complex rather than a separate subunit that is recruited on target DNA binding. Although the Cas3 structure was not observed in the cryo-EM density, bioinformatic predictions suggest that the domain organisation is distinct from Cas3s from other type I CRISPR subtypes, with its HD nuclease domain located at the C-terminus instead of the N-terminus (7) (**Figure 1)**. By comparison with the crystal structure of Cas3 from *Thermobifida fusca*, we could predict the identity of key active site residues in the HD nuclease and helicase domains of the *T. sulfidiphilus* Cas3 protein **(Figure 1)**.

**Figure 1.**
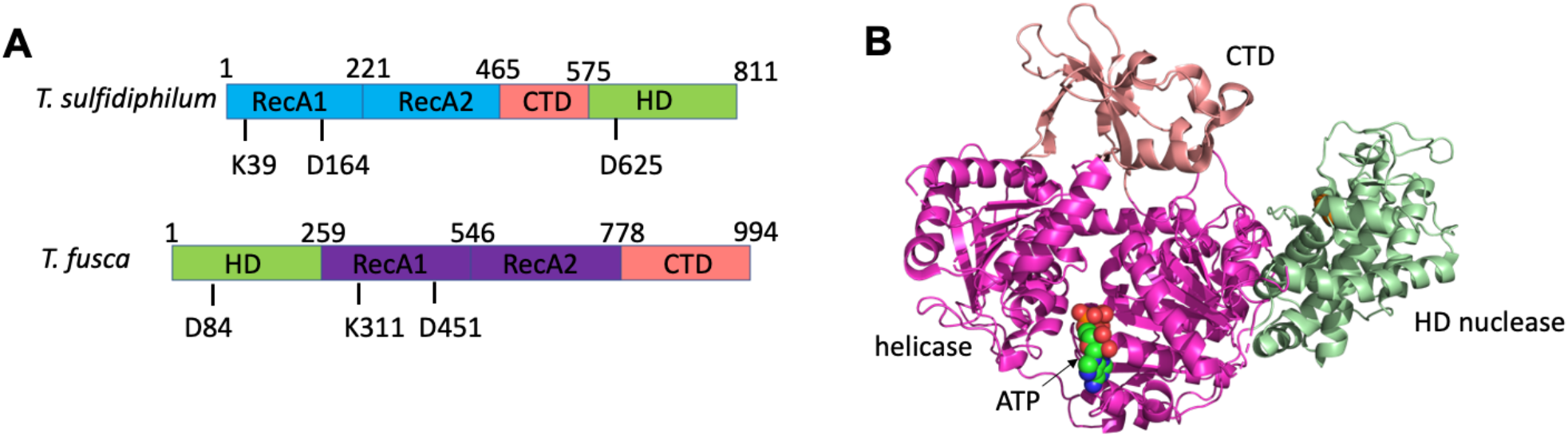
Comparison of *T. fusca* and *T. sulfidiphilus* Cas3. **A**. Domain organisation and active site residues of *T. fusca* and *T. sulfidiphilus* Cas3. Each has two RecA-family motor domains containing active site residues that define the Walker A and Walker B boxes of the helicase motor. The HD nuclease domain is present at the N-terminus of *T. fusca* Cas3 but at the C-terminus of *T. sulfidiphilus* Cas3. Each has several conserved acidic residues that coordinate the catalytic metal ions in the active site, of which D625/D84 is labelled. **B**. X-ray crystal structure of *T. fusca* Cas3 bound to the ATP analogue AMP-PNP (35), coloured to match the domains shown in A.

### The importance of Cas3 helicase and nuclease activities for defence against MGEs

Cas3 has two active sites – a 3’-5’ SF2 helicase and an HD nuclease (**Figure 1**). To dissect the importance of these two activities for defence against MGE, we constructed three variant Cas3 proteins: one with a K39A mutation of the key Walker A motif, one with a D164A mutation of the Walker B motif and a third with a D625A mutation in the active site of the HD domain. Variant proteins equivalent to D164A and D625A were previously explored for *T. fusca* Cas3 (35). We proceeded to investigate the phenotypes of these variants using our established plasmid challenge assay (39) (**Figure 2A**). In this assay, cells with a functional *T. sulfidiphilus* Cascade were challenged with a plasmid containing a DNA sequence targeted by the crRNA. If the targeted plasmid is destroyed, cells do not become resistant to tetracycline and no colonies are observed. Consistent with previous studies, cells with wild-type Cas3 were fully resistant to plasmid challenge, regardless of whether full gene expression of Cascade was induced by lactose and arabinose (**Figure 2A**). In contrast, cells with the K39A and D164A variant of Cas3, which lack helicase activity, were only resistant when expression was fully induced. These data suggest that when Cas3 cannot translocate on DNA, but can still cut it, type I-G CRISPR defence is weakened but not abolished.

**Figure 2.**
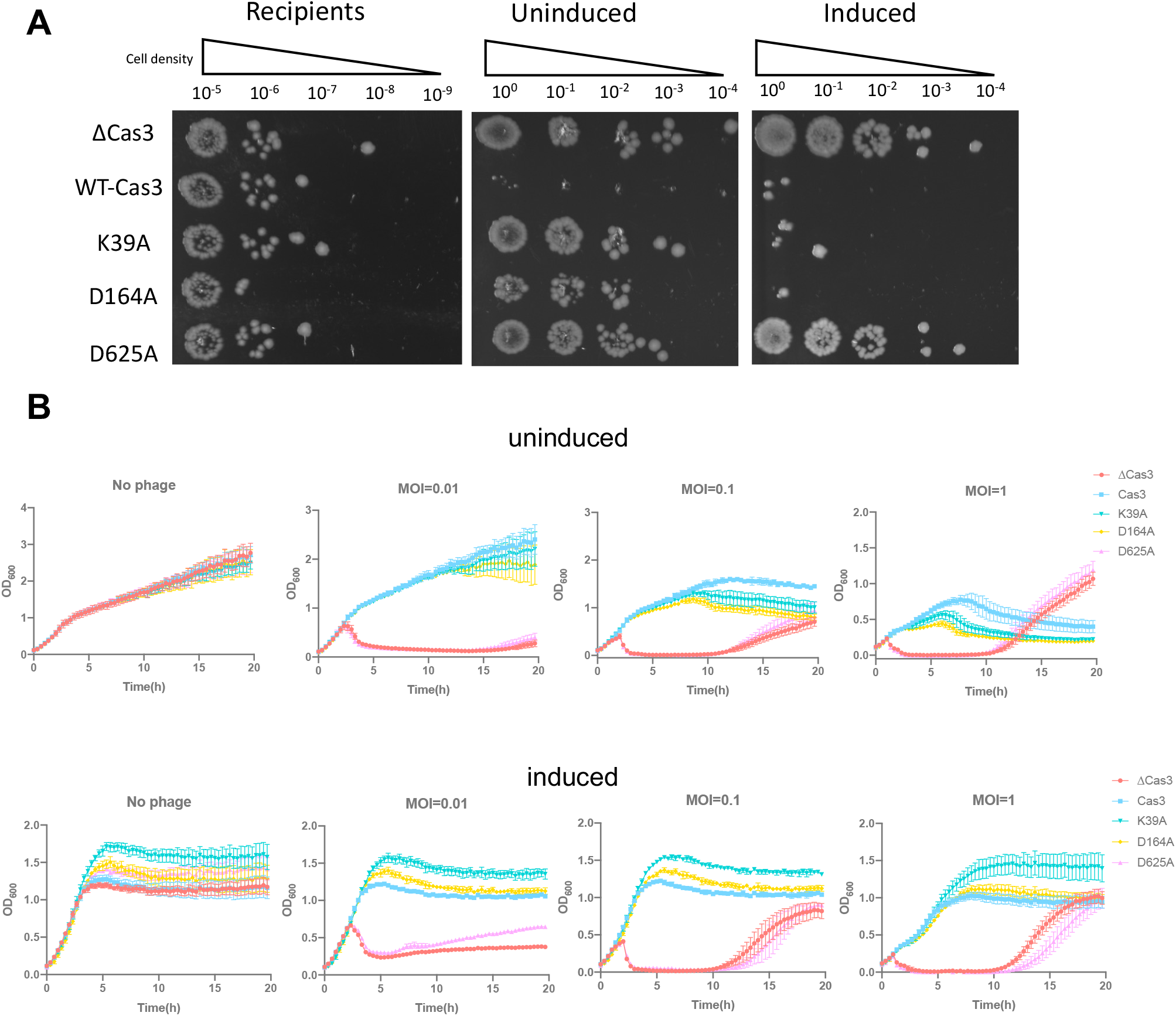
Plasmid and Phage P1 Challenge with wild-type and variant Cas3. **(A)** Cells on the plates in different condition. Recipients, Ampicillin and Spectinomycin in plates; Uninduced, Ampicillin, Spectinomycin and Tetracycline in plates; Induced, with all three antibiotics and lactose, arabinose for induction of the type I-G CRISPR system. The Cas3 variants ΔCas3, K39A, D164A and D625A were also studied. **(B)** Cell growth curve under phage P1 challenge (MOI=0, 0.01, 0.1 and 1). ΔCas3 cells lack the *cas3* gene. Data points represent the mean of six experimental replicates (two biological replicates and three technical replicates) with standard deviation shown.

When submitted to a phage P1 challenge assay, cells expressing the K39A or D164A Cas3 variant showed full immunity against phage infection under induced condition, and compromised immunity at higher MOIs compared to wild-type Cas3 when expression was uninduced (**Figure 2B**). In previous *in vitro* experiments, we observed that *T. sulfidiphilus* Cascade in the absence of ATP (and thus helicase activity) only cleaved target DNA at the site of Cas3 loading (39). This appears to be enough *in vivo* to prevent target plasmid and phage replication as long as the type I-G CRISPR effector is highly expressed or MOI is low. In marked contrast, the D625A variant of Cas3 was indistinguishable from the ΔCas3 strain (**Figure 2A&2B**). To investigate this further, we expressed and purified the Cas3 D625A variant from *E. coli*. Unlike the wild-type protein, the D625A variant eluted as a large aggregate from a size exclusion column (**Figure S1**). These data suggest that mutations disrupting the iron binding site of the Cas3 nuclease domain may disrupt protein folding rather than just inactivating the nuclease domain. This observation emphasises the importance of checking the phenotypes of variant proteins *in vitro* as well as *in vivo*.

### Genome engineering with the Type I-G CRISPR system

We proceeded to explore the potential of the type I-G system in genome engineering. We first constructed the vector pM2, containing the type I-G operon under arabinose-inducible promoter control (**Figure 3**). pM2 was transformed into *E. coli* MG1655 along with plasmid pSPACER to produce pre-crRNA for *lacZ* targeting. Cells were spread on X-gal plates for blue-white screening (**Figure 3C**). In the absence of a targeting crRNA, large numbers of blue cells were observed. When a spacer targeting the *lacZ* gene on the bacterial genome was induced, colony counts were significantly reduced and both blue and white colonies were observed (**Figure 3D**).

**Figure 3.**
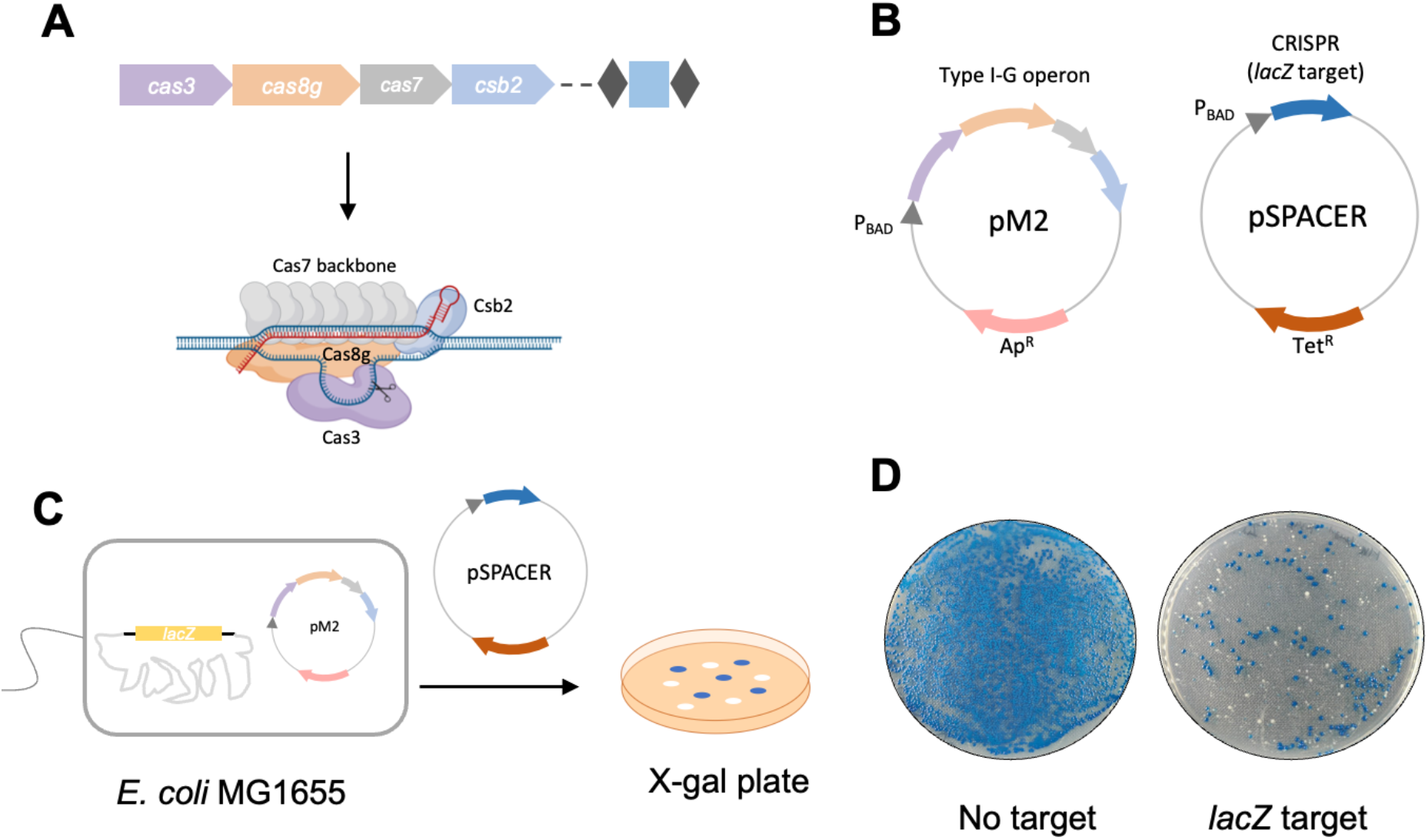
Target editing by type I-G. **(A)** An overview of type I-G CRISPR operon from *T. sulfidiphilus* HL-EbGr7. *cas* proteins (7 x Cas7, Csb2, Cas8g and Cas3) together with crRNA form the effector complex and target dsDNA. **(B)** pM2 vector contains type I-G operon (*cas3, cas8g, cas7* and *csb2*) under arabinose promoter control; pSPACER vector, with CRISPR repeat and spacer (target *lacZ*) under arabinose promoter control. **(C)** For *lacZ* disruption, pM2 was transformed into *E. coli* MG1655 strain *(lacZ* intact) first, and pSPACER was introduced to activate type I-G *lacZ* targeting, cells were spread on the X-gal plates for blue-white screening. **(D)** A representative blue-white screening assay with no target control and *lacZ* target spacer.

When we first introduced the type I-G system into *E. coli* for self-genome targeting, we noticed a significant decrease (around 2 orders of magnitude) in cell number compared to empty vector (no target spacer) control (**Figure 4A**). Even in the absence of arabinose induction, a loss of cells was still observed, suggesting that the system was actively targeting the *lacZ* gene. On blue/white screening, white colonies only appeared on the plates with a targeting spacer (**Figure 4B**). The reduced cell counts resulting from experiments targeting the *lacZ* gene were most likely due to the generation of unrepaired double strand breaks (DSB) and/or deletion of an essential flanking gene. Similar toxicity upon self-targeting has been seen in endogenous type I CRISPR and heterologous Cas9 chromosomal targeting in prokaryotic cells (16, 21, 44). In particular, Cas3 of other type I CRISPR systems is responsible for long-range deletions of chromosomal DNA (14, 30).

**Figure 4.**
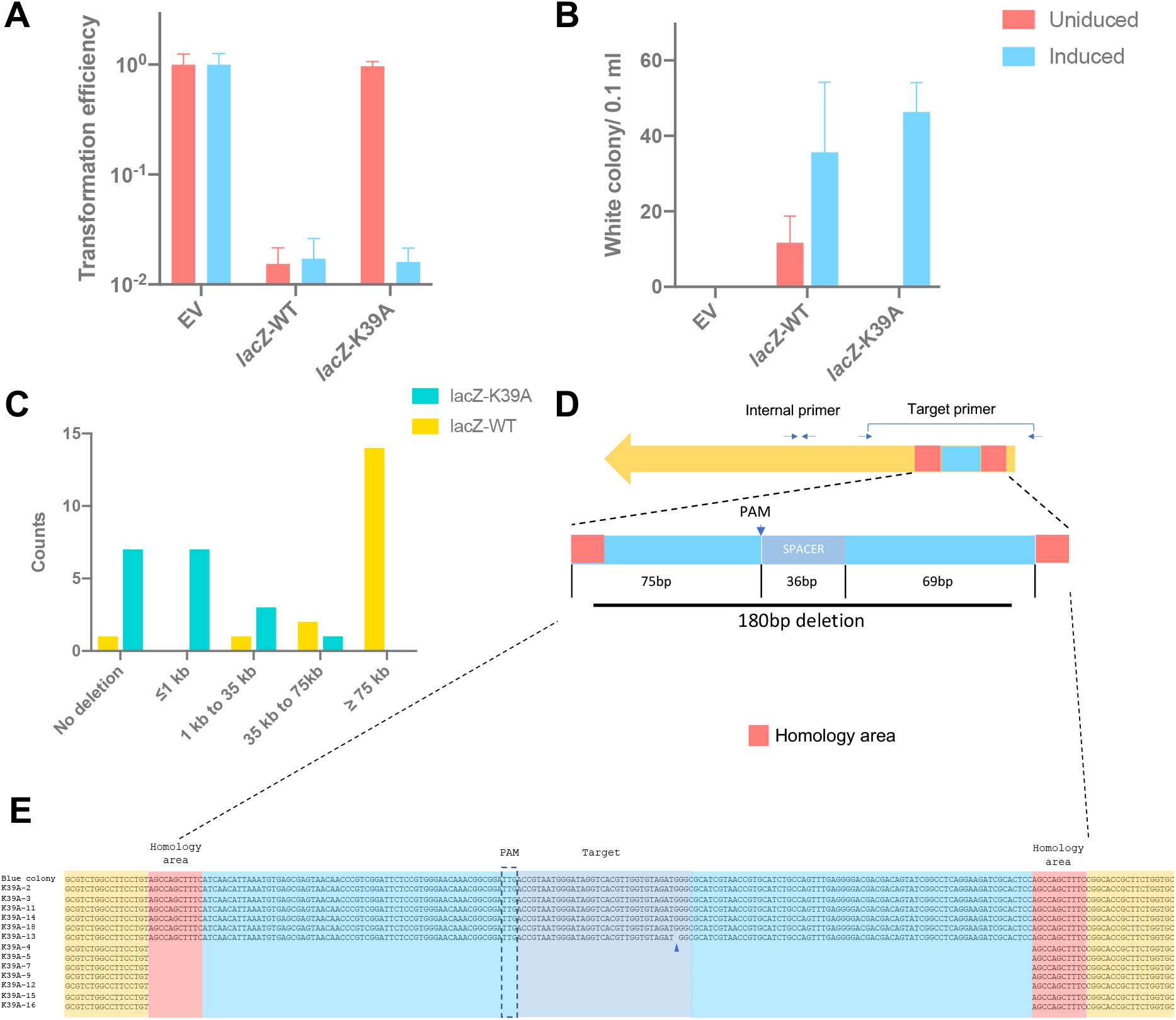
Type I-G editing on *lacZ*. **(A, B)** Transformation efficiency and white colony number on the plate after transformation of the empty vector control or *lacZ* target spacer to wildtype Cas3 strain or K39A Cas3 strain with L-arabinose induction (blue) or without induction (red); Transformation efficiency was calculated as the number of transformants divided by the number of transformants for original plasmid without target (Empty vector control); Values and error bars represent the mean of three biological replicates and standard deviation. **(C)** Editing outcome of *lacZ* targeting by wildtype type I-G (yellow) or Cas3 K39A (green); 18 white colonies generated by *lacZ* targeting were submitted for tiling PCR to determine the deletion range; counts of white colonies are plotted against deletion range. **(D)** A schematic of 180 bp deletion by Cas3 K39A targeting *lacZ*; Blue, deleted sequence; Red, homology area; PAM, protospacer adjacent motif; SPACER, spacer sequence for *lacZ* targeting; Primers used for amplification of target area were indicated by arrows. **(E)** Sequence analysis of the 180 bp deletion by Cascade with Cas3 K39A; Blue arrow, a point mutation was detected; Direct repeats in the flanking sequence are shown in red.

Given the large decrease in viable cells and the extensive genomic deletions observed in survivors, we next explored *lacZ* gene disruption using the Cas3 K39A variant. Cells expressing the K39A variant of Cas3 as part of *T. sulfidiphilus* Cascade complex did not experience cell death in the absence of arabinose induction (**Figure 4A**), but when fully induced we observed roughly two logs reduction in viable cells, similar to cells with wild-type Cas3 (**Figure 4A**), with survivors displaying a *ΔlacZ* phenotype in similar numbers to the wild-type system (**Figure 4B**).

To test whether type I-G Cas3 yielded long-range deletion on genome targeting, white colony survivors were submitted for tiling PCR. 17 out of 18 white survivors from induced plates had a *lacZ* target deletion (**Figure 4C, S2, S3**). Out of 18 white colonies, 13 survivors had a deletion at least as far as 55 kb downstream of *lacZ*, 3 survivors experienced a deletion of 45 kb to 55 kb downstream and 1 survivor had a shorter 10 kb to 25 kb downstream deletion. The *lacZ* gene of one white colony (WT-18) remained intact with no detected deletion around the target site, suggestive of a spontaneous *lacZ* mutation. The essential *hemB* gene is located 20 kb upstream of *lacZ*. Unsurprisingly, all the survivors had an intact *hemB* gene locus, but 17 colonies showed 10-20 kb deletions of upstream DNA. In subsequent experiments, we targeted the *lacZ* adjacent genes *yahK* and *frmA*. When the downstream gene *yahK* was targeted, the yield of white survivors was comparable to *lacZ* target, while the upstream gene *frmA* target sharply lowered the number of white surviving colonies obtained, likely due to its proximity to the essential gene *hemB* (**Figure S4**). These data suggest that the type I-G system yields long-range bidirectional degradation of the *E. coli* genome,

In contrast, PCR tiling revealed a marked difference in editing outcomes when using the *cas3* K39A mutant. Out of 18 white colony survivors analysed by tiling PCR, 14 gave a PCR product when using an internal *lacZ* primer, 4 had a localised *lacZ* deletion, and only 1 of them showed long-range deletion (**Figure 4C, 4D, S3, S5**). These white survivors were further analysed using PCR primers that covered the target area.

Intriguingly, PCR products from these survivors varied in size, indicative of small deletions **(Figure S5C)**. Out of 13 sequenced PCR products, 1 contained a point mutation, 7 had a precise 180 bp deletion flanking the target site and the other 5 had no obvious edit (**Figure 4E**). Close investigation of the sequences revealed the presence of 11 bp direct repeats flanking the deleted region (**Figure 4E**). This suggests that the DNA break introduced by Cas3 was repaired by limited strand resection and repair by microhomology-mediated end joining (MMEJ). Gene deletions flanked by short regions of microhomology were recently observed in a study of genome engineering in *Bifidobacteria* (24).

To investigate this phenomenon in more detail, we targeted a second site in the *lacZ* gene using a different crRNA with the K39A Cas3 variant Cascade and analysed it as before. DNA sequencing showed various editing outcomes: intact target site (8 out of 17 tested colonies), point mutation (2/17), 24 bp short deletion (5/17), 324 bp long deletion (1/17) and a 11 bp insertion (1/17) (**Figure S6**). Microhomologies ranging from 2 to 8 bp were apparent flanking the deleted regions (**Figure S6**). Overall, genome targeting via Cas3 variant K39A generated distinct editing outcomes compared with the wild-type editing.

### Utilising the type I-G system in homology-directed repair

In the absence of any donor DNA to direct repair, the main products of gene disruption likely arise from error-prone end joining pathways. We proceeded to investigate whether the provision of donor DNA as a homology-directed repair (HDR) template would result in different outcomes. To investigate this, the pHR vector was constructed with a donor sequence incorporated downstream of the CRISPR array (**Figure 5A&S7A**). If successful, the design of the HDR donor should result in a 13 kb deletion in the genome. For the type I-G system with wildtype Cas3, even with homologous recombination templates, the survivability was still low and target HDR efficiency was under 20 % (1 target deletion out of 8 assayed white colonies) (**Figure 5**). However, when targeting with Cas3 K39A, an increase in cell survivability and white colony number was observed. The target editing efficiency was over 90 % (28 target deletions out of 30 white colonies) (**Figure 5D**) and a 13 kb deletion was observed (**Figure S7**). Overall, the introduction of donor DNA for HDR increased the cell survivability and yielded the desired long-range deletion in the bacterial genome when the K39A variant was used.

**Figure 5.**
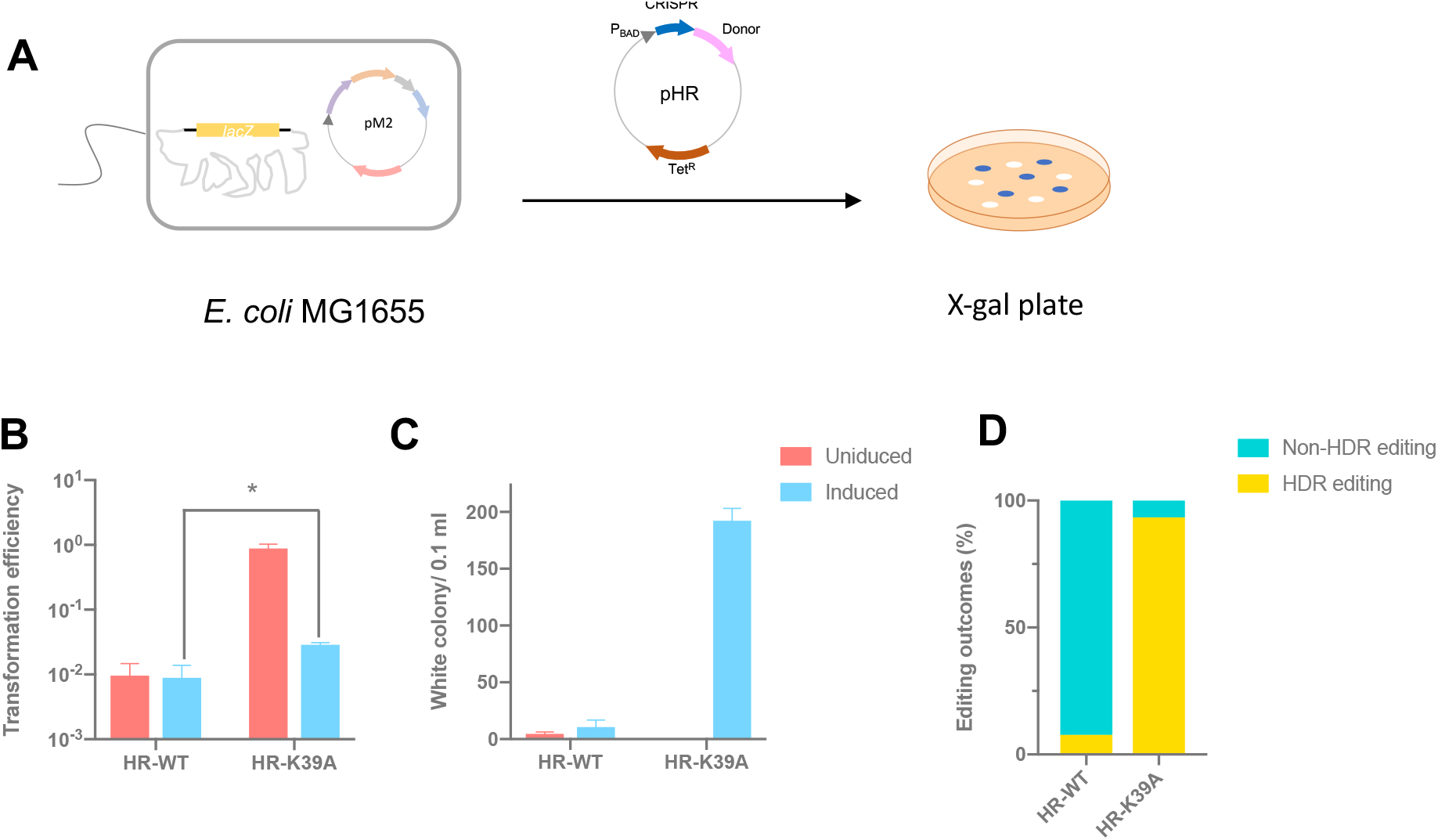
Homology-directed Repair. **(A)** A schematic of homology directed repair; pM2 was transformed into *E. coli* MG1655 strain *(lacZ* intact) first, and pHR with donor sequence and *lacZ* target spacer was introduced to activate type I-G *lacZ* targeting and desired HDR, cells were spread on the X-gal plates for blue-white screening. **(B, C)** Transformation efficiency and white colony number on the plate after transformation of pHR into WT *cas3* strain or K39A *cas3* mutant strain with L-arabinose induction (blue) or without induction (red); Transformation efficiency was calculated as the number of transformants divided by the number of transformants for original plasmid without target (Empty vector control). Values and error bars represent the mean of three biological replicates and standard deviation; * p<0.05, paired T-test. **(D)** Editing outcomes with donor templates; Percentage of white colonies with desired HDR editing (yellow), without HDR editing (green); 30 individual white colonies from HR-K39A, 13 from HR-WT were assayed.

## Discussion

Although Cas3 translocates unidirectionally on target DNA *in vitro* (34, 37), type I CRISPR systems are capable of generating both unidirectional (27, 28, 30, 31, 32, 33) and bi-directional (14, 26, 29) deletions in target genomes. Here, we have shown that type I-G Cascade-Cas3 specifically creates bi-directional long-range genome deletions in *E. coli*. Similar bidirectional degradation has been observed for the type I-C CRISPR in *Pseudomonas aeruginosa* (14), where an average deletion of around 90 kb was observed. However, when the type I-C system from *Neisseria lactamica* was repurposed for application in eukaryotes, it resulted in unidirectional degradation (28). These observations suggest that editing outcomes are dependent on both the specific type I CRISPR system and the experimental species under study.

Cas3 is universally conserved across all type I CRISPR systems, despite the differences in Cascade composition (7). In the type I-G system Cas3 is an integral subunit of Cascade rather than being recruited on target DNA binding (24, 39). This association between Cas3 and Cascade has also been observed in a type I-A system where the authors proposed an allosteric activation mode, in contrast to the common trans-recruitment mode for Cas3 (26, 43). The type I-G Cas3 shares common features with canonical Cas3 (a SF2-helicase domain and a HD-nuclease domain), but at the protein sequence level the HD nuclease domain is found at the C-terminus instead of N-terminus. In type type I-A systems, Cas3 is encoded on two individual genes that sepately express the helicase and HD nuclease domains (7). These differences in Cas3 domain organisation seem to correlate with the existence of two different Cascade-Cas3 activation modes.

While the type I-G system with wild-type Cas3 generates large bi-directional deletions, we have shown that the helicase activity of Cas3 is not essential for functional immunity against MGEs in *E. coli*, as long as the proteins are well expressed. This suggests that localised nicking of the target strand in targeted MGEs can be sufficient to prevent replication, possibly because replication forks collapse when encountering a nick in the leading strand template, resulting in a double strand break. Likewise, the use of the helicase-deficient K39A variant of Cas3 did not abolish gene disruption in *E. coli*. However, the edited products were fundamentally different, with small deletions predominating. The recurrence of a specific 180 bp deletion was initially surprising, but may be explained by the presence of direct repeats flanking the deleted sequence, which could allow end joining following limited strand resection. Microhomology mediated end joining is observed when HDR is not an option in *E. coli* (45), and has been seen previously for Cas3-mediated deletions in *P. aeruginosa* (14).

Provision of a donor template to enhance HDR during edits resulted in the expected HDR outcomes. In particular, the use of the Cas3 K39A variant in combination with a donor template enhanced editing efficiency. Recently, a Cas3 helicase variant of the *Zymomonas mobilis* type I-F system was applied endogenously to carry out genome editing with high efficiency (46). In that study, crRNA was used to target each strand of the target, resulting in “dual nicking” of the target gene. Our data suggest that targeting genes with a single guide RNA can still lead to efficient gene disruption and gene replacement, simplifying the procedure.

For obvious reasons, the compact Cas9 and Cas12 enzymes have been favoured for genome editing in a wide range of species. Nonetheless, type I systems have now been widely used to create larger edits in a range of cognate and heterologous cell types. Here, we have demonstrated that the relatively simple, 4-gene type I-G system can be used to generated gene deletions and replacements in *E. coli*. Surprisingly, abolishing the helicase activity of Cas3 does not prevent either effective immunity against MGE, and can result in enhanced genome editing, suggesting that the cellular responses to the DNA nicks produced can result in gene disruption with smaller deletions, while still supporting HDR when a doner template is provided.

**Table 1.**
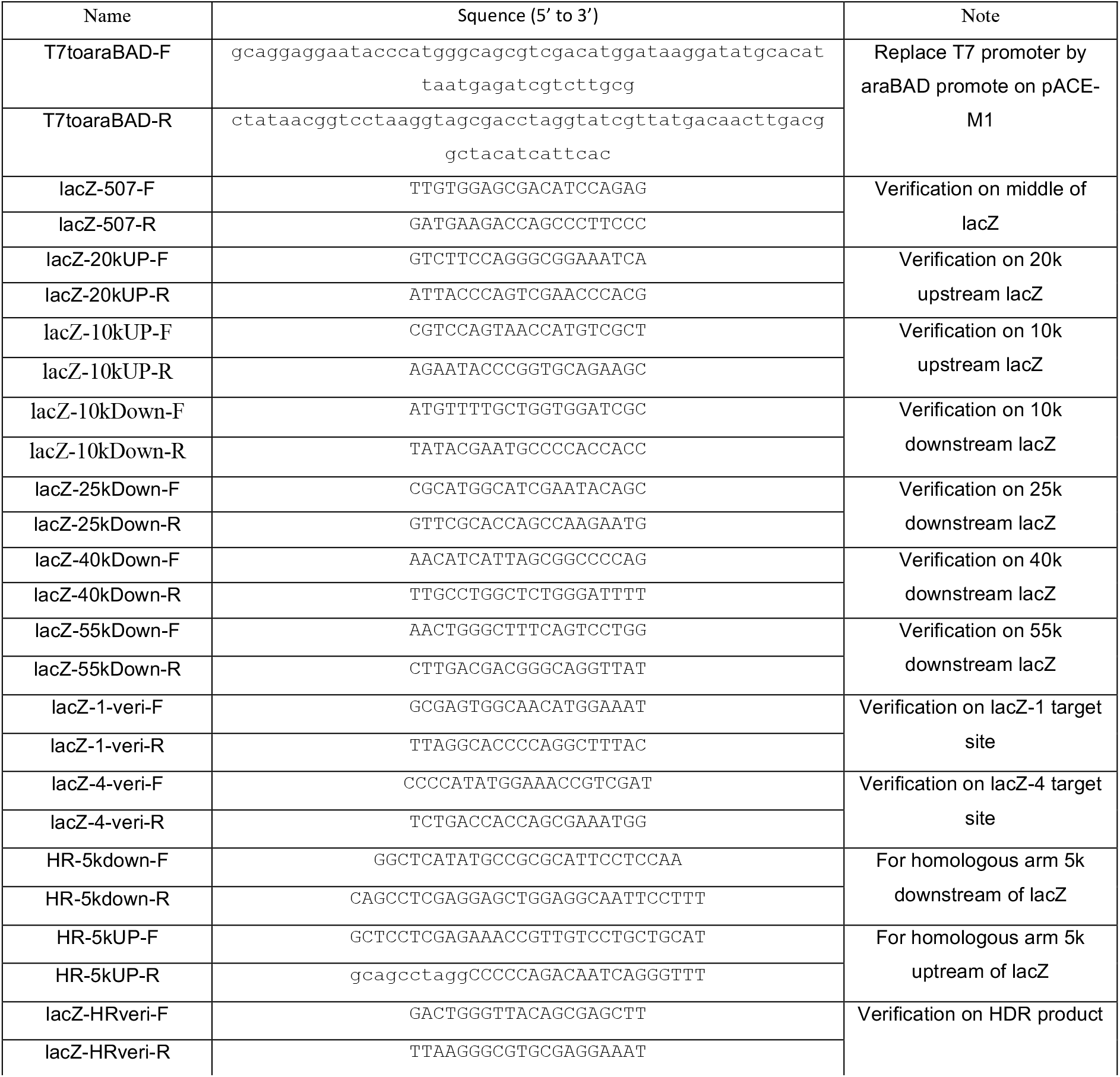
Primers.

**Table 2.**
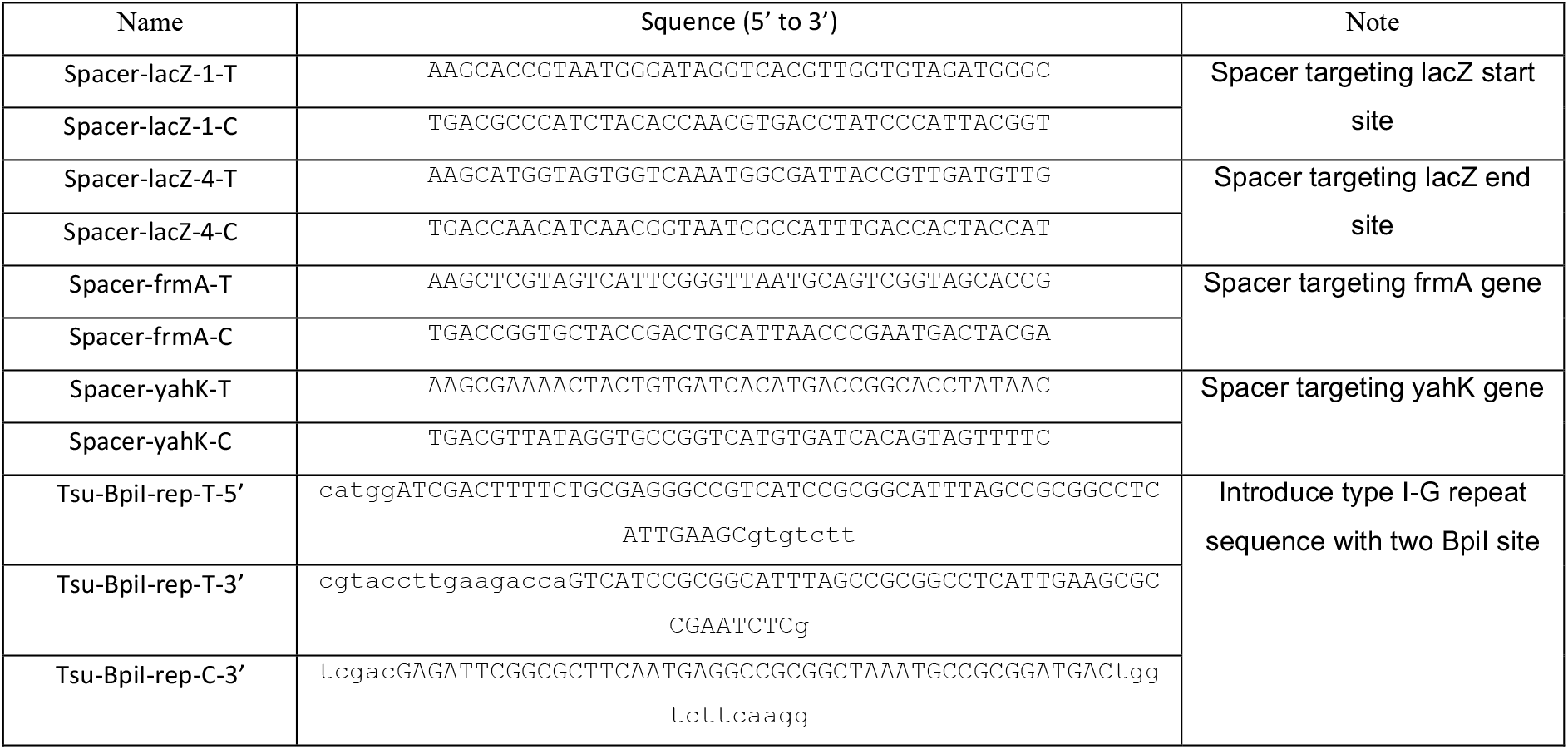
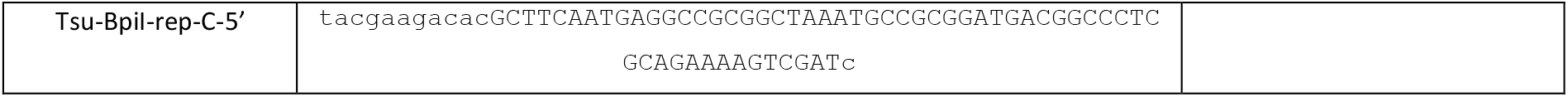
oligonucleotides.

**Table 3.** Colony counts.

| Figure 4A CFU/0.1 ml in log |  |  |  |  |  |  | Figure 4B white colony number/0.1 ml |  |  |  |  |  |
| --- | --- | --- | --- | --- | --- | --- | --- | --- | --- | --- | --- | --- |
|  | Uniduced |  |  | Induced |  |  | Uniduced |  |  | Induced |  |  |
| EV | 5.26 | 5.30 | 5.08 | 5.16 | 5.27 | 5.05 | 0.00 | 0.00 | 0.00 | 0.00 | 0.00 | 0.00 |
| lacZ-WT | 3.57 | 3.37 | 3.22 | 3.60 | 3.34 | 3.15 | 18.00 | 13.00 | 4.00 | 51.00 | 41.00 | 15.00 |
| Figure 4A CFU/0.1ml in log |  |  |  |  |  |  | Figure 4B white colony number/0.1 ml |  |  |  |  |  |
|  | Uniduced |  |  | Induced |  |  | Uniduced |  |  | Induced |  |  |
| EV | 5.35 | 5.33 | 5.31 | 5.30 | 5.37 | 5.28 | 0.00 | 0.00 | 0.00 | 0.00 | 0.00 | 0.00 |
| lacZ-K39A | 5.36 | 5.30 | 5.27 | 3.59 | 3.61 | 3.31 | 0.00 | 0.00 | 0.00 | 40.00 | 55.00 | 44.00 |
| Figure 5B CFU/0.1ml in log |  |  |  |  |  |  | Figure 5C white colony number/0.1 ml |  |  |  |  |  |
|  | Uniduced |  |  | Induced |  |  | Uniduced |  |  | Induced |  |  |
| HR-WT | 3.41 | 3.13 | 2.95 | 3.33 | 3.01 | 2.89 | 5.00 | 6.00 | 3.00 | 16.00 | 12.00 | 4.00 |
| HR-K39A | 5.21 | 5.35 | 5.25 | 3.81 | 3.76 | 3.75 | 0.00 | 0.00 | 0.00 | 200.00 | 180.00 | 197.00 |
| Figure S2A CFU/0.1ml in log |  |  |  |  |  |  | Figure S2B white colony number/0.1 ml |  |  |  |  |  |
|  | Uniduced |  |  | Induced |  |  | Uniduced |  |  | Induced |  |  |
| EV | 5.26 | 5.30 | 5.08 | 5.16 | 5.27 | 5.05 | 0.00 | 0.00 | 0.00 | 0.00 | 0.00 | 0.00 |
| lacZ | 3.57 | 3.37 | 3.22 | 3.60 | 3.34 | 3.15 | 18.00 | 13.00 | 5.00 | 51.00 | 41.00 | 12.00 |
| frmA | 3.40 | 3.40 | 3.18 | 3.40 | 3.36 | 3.05 | 1.00 | 0.00 | 0.00 | 3.00 | 1.00 | 0.00 |
| yahK | 3.35 | 3.33 | 3.25 | 3.39 | 3.30 | 3.14 | 26.00 | 12.00 | 8.00 | 44.00 | 33.00 | 9.00 |

## Conflicts of Interest

The authors declare they are aware of no conflicts of interest.

## Funding information

This work was supported by the Biotechnology and Biological Sciences Research Council (REF: BB/S000313/1 to MFW), and the China Scholarship Council (REF: 202008060345 to QS).

---

**FigureS1.**
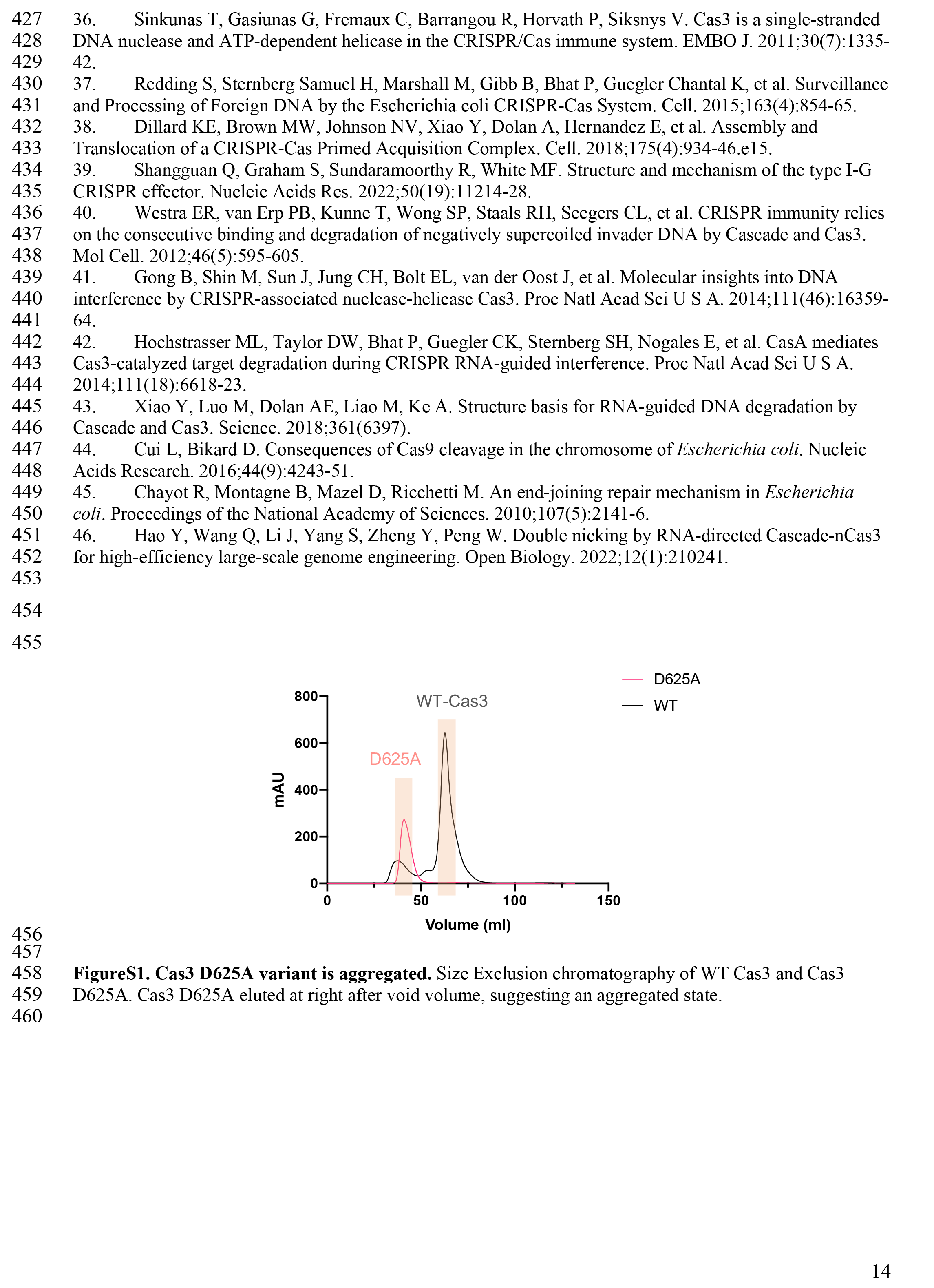
Cas3 D625A variant is aggregated. Size Exclusion chromatography of WT Cas3 and Cas3 D625A. Cas3 D625A eluted at right after void volume, suggesting an aggregated state.

**FigureS2.**
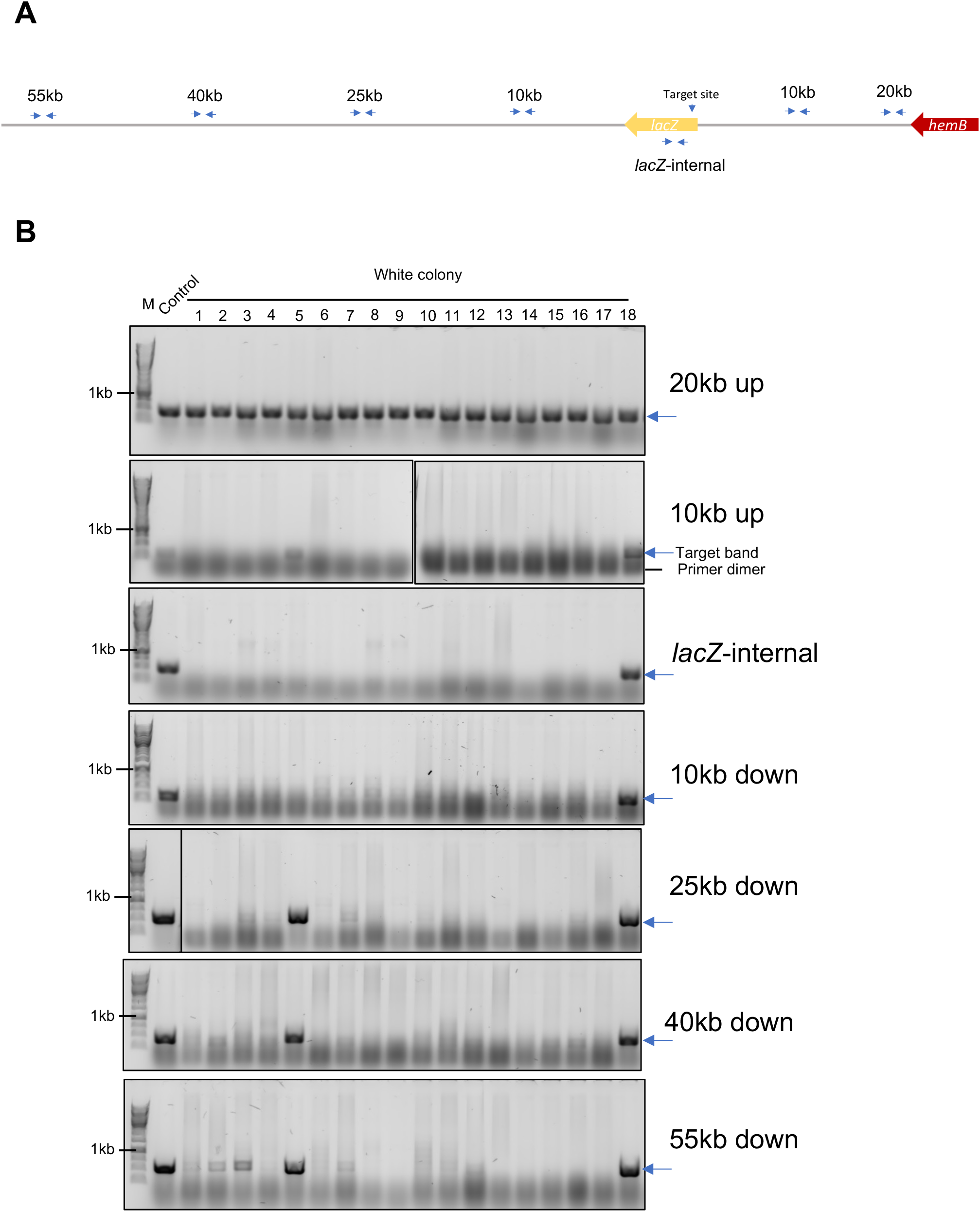
Tiling PCR of type I-G targeting *lacZ*. **(A)** An overview of location of tiling PCR primers; A pair of small blue arrow represents a set of PCR primers. **(B)** Tiling PCR product from different primers was submitted for electrophoresis on a 0.8% agarose gel. 18 white colonies and 1 blue colony (Control) were assayed. M, marker. Blue arrows indicate positive PCR products.

**Figure S3.**
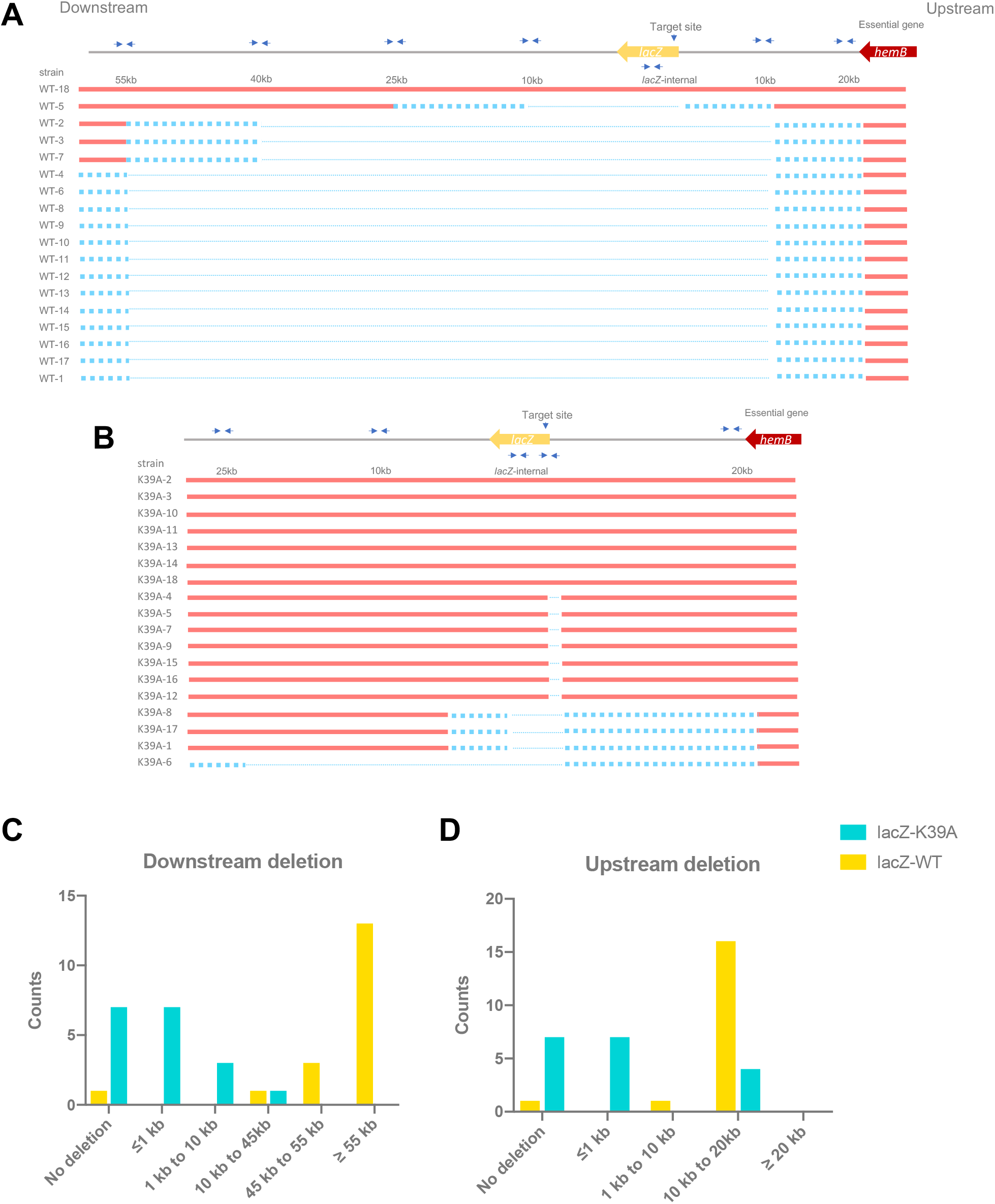
Type I-G editing on *lacZ*. **(A) (B)** A deletion map showing the outcome of *lacZ* targeting by wildtype type I-G or Cas3 K39A; 18 white colonies generated by *lacZ* targeting were submitted for tiling PCR to determine the deletion range; Pairs of small blue arrow represent tiling PCR primers; Red lines indicate the intact sequence on genome; Blue dot lines indicates possible deleted sequence; Thin blue dash lines indicate confirmed deleted sequence. **(C) (D)** Editing outcome of *lacZ* targeting by wildtype type I-G (yellow) or Cas3 K39A (green); 18 white colonies generated by *lacZ* targeting were submitted for tiling PCR to determine the deletion range; Counts of white colonies are plotted against deletion range.

**FigureS4.**
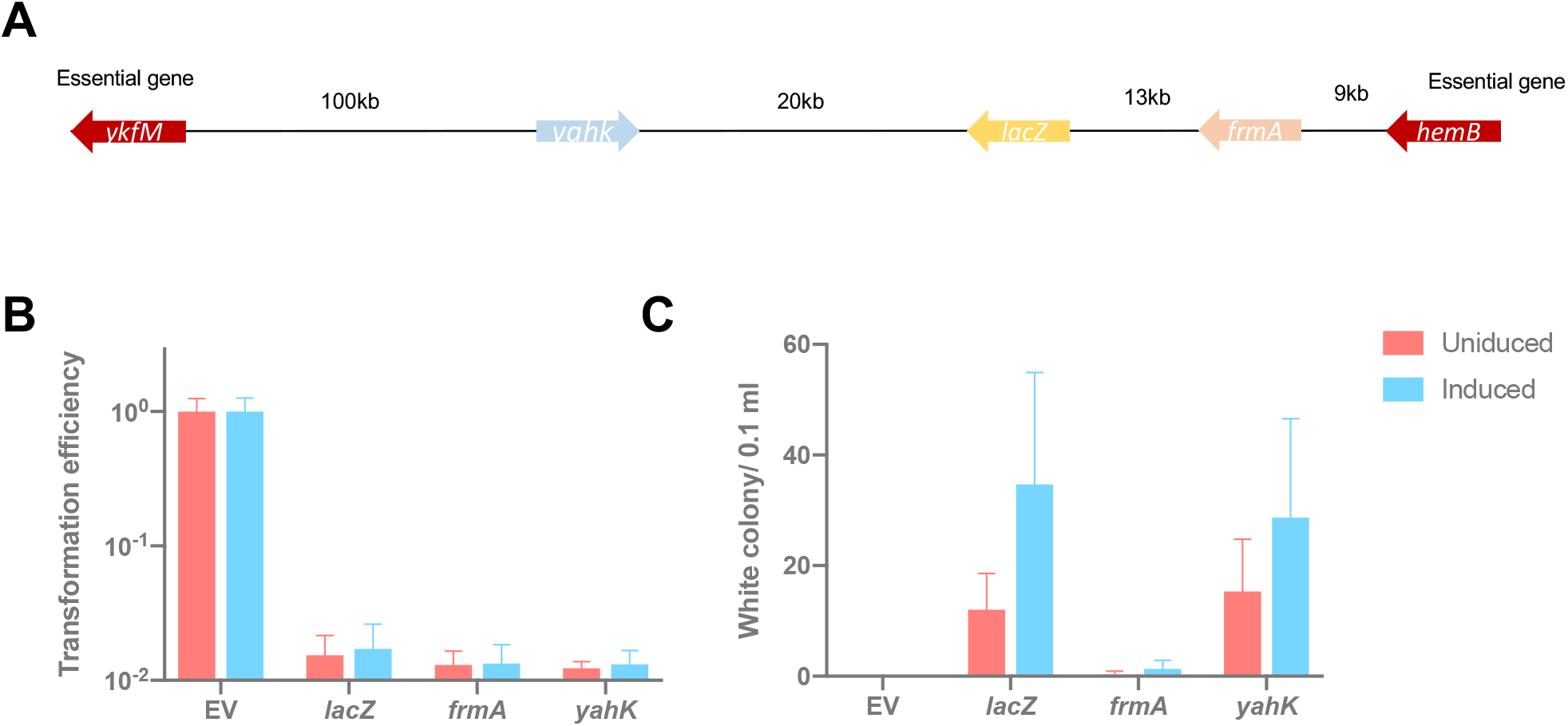
*yahK* and *frmA* target editing. **(A)** *yahK* and *frmA* location on *E*.*coli* genome. **(B) (C)** Transformation efficiency and white colony number on the plate after transformation of the empty vector, *lacZ* target spacer, *frmA* target spacer or *yahK* target spacer respectively with L-arabinose induction (blue) or without induction (red); Transformation efficiency was calculated as the number of transformants divided by the number of transformants for original plasmid without target (Empty vector control); Values and error bars represent the mean of three biological replicates and standard deviation.

**FigureS5.**
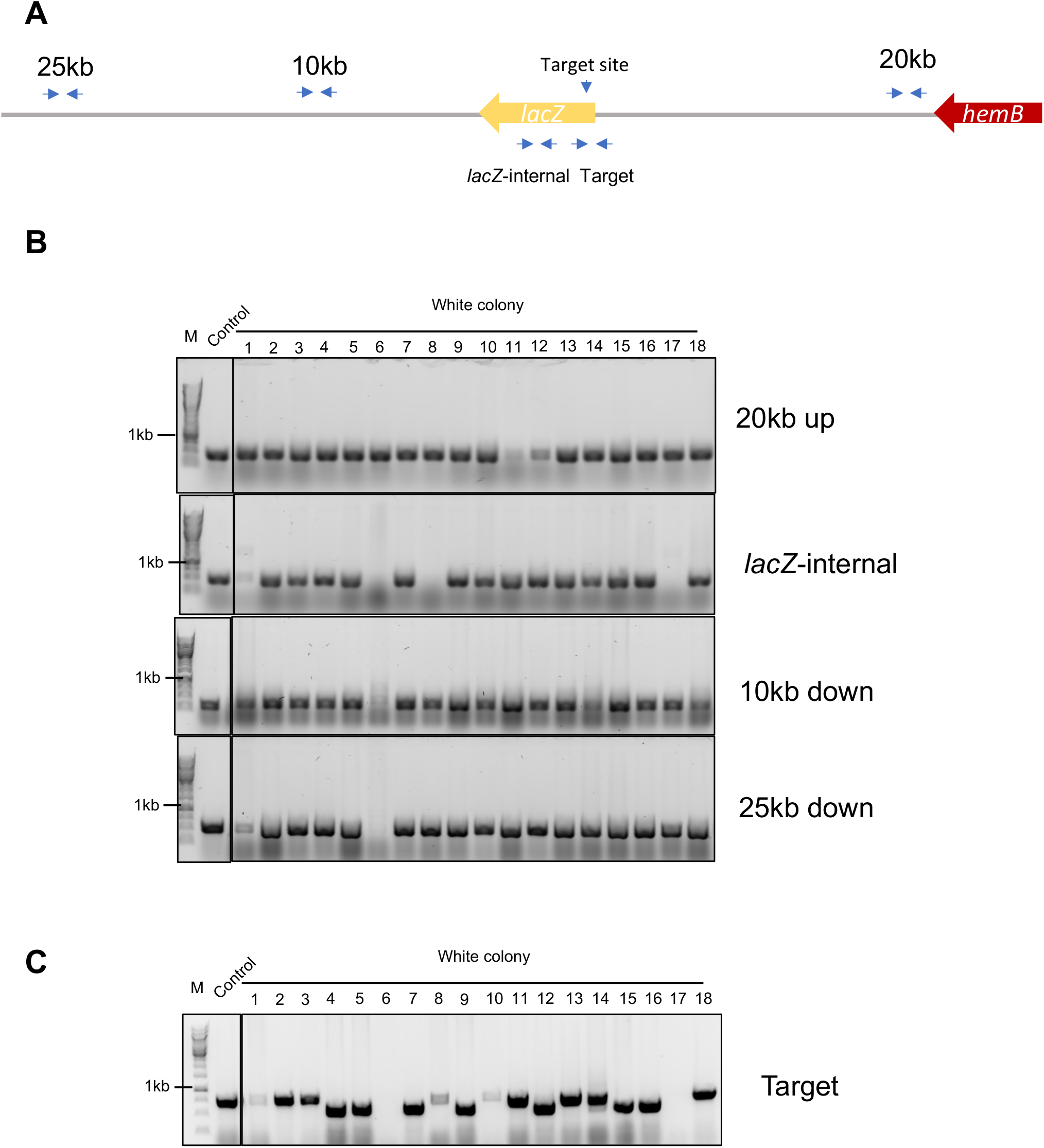
Tiling PCR of type I-G K39A targeting *lacZ*. **(A)** An overview of location of tiling PCR primers; A pair of small blue arrow represents a set of PCR primers. **(B)** Tiling PCR product from different primers was submitted for electrophoresis on a 0.8% agarose gel. 18 white colonies and 1 blue colony (Control) were assayed. M, marker. **(C)** PCR product with primers that cover the target area was submitted for electrophoresis on a 0.8% agarose gel.

**FigureS6.**
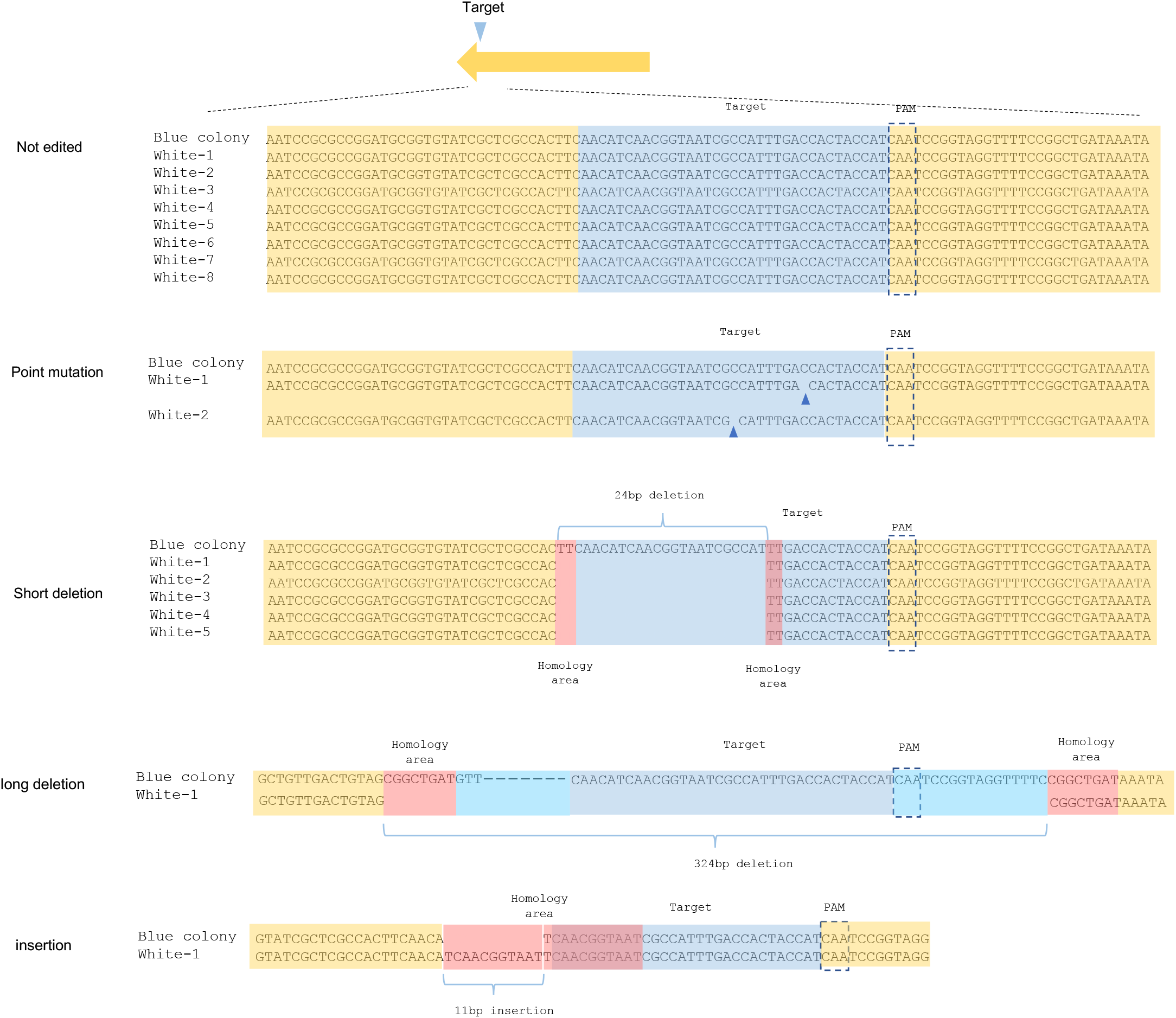
K39A target deletion on alternative *lacZ* site. A different target site on *lacZ* gene produces various editing outcomes; Blue arrow, point mutation site. Homology in red.

**FigureS7.**
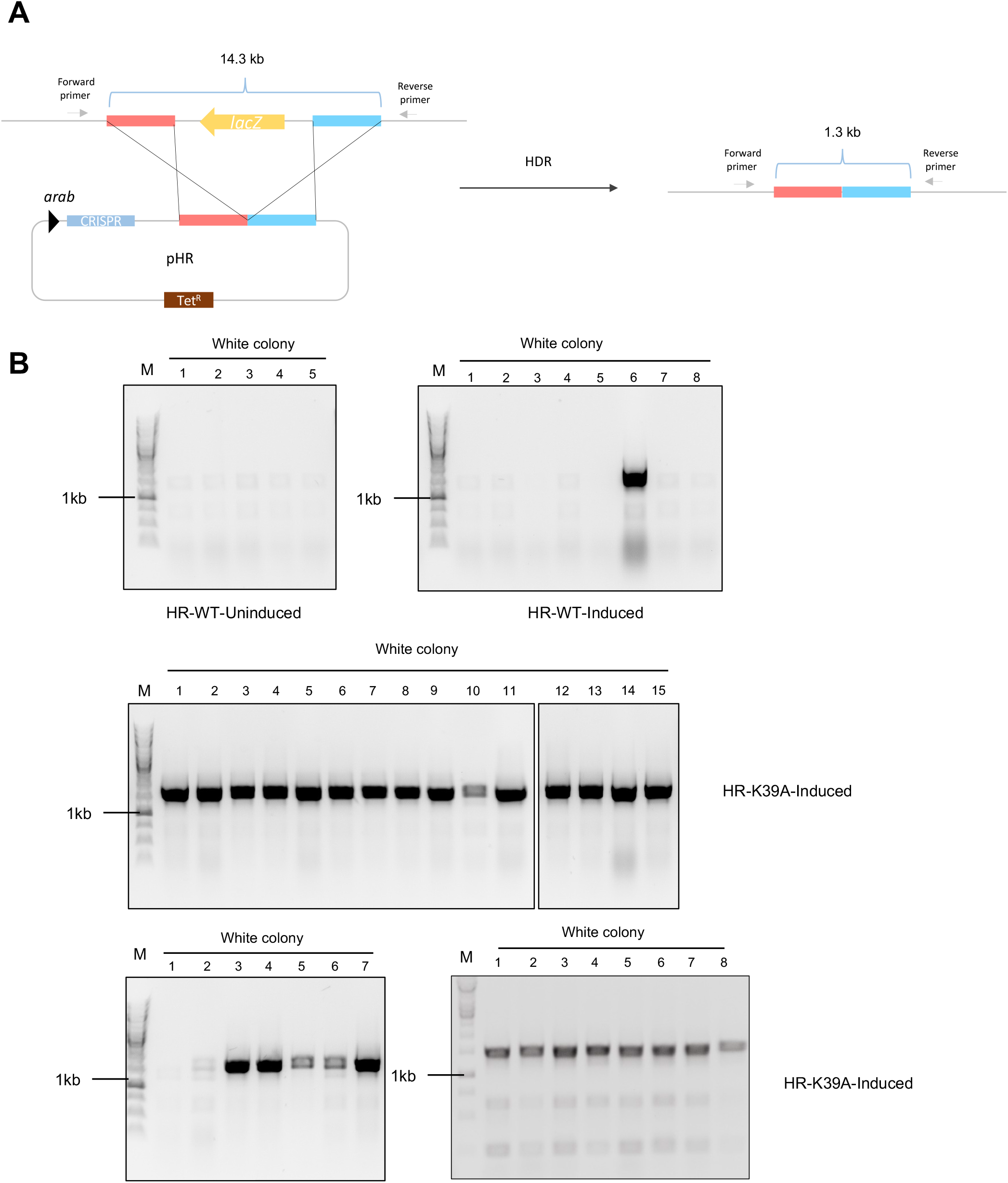
Homologous recombination for repair. **(A)** A schematic of homologous recombination; Homologous arms in red and blue; Primers for desired HDR verification was shown in grey arrow. **(B)** PCR product with verification primers were submitted for electrophoresis on a 0.8% agarose gel; 1.3kb product was detected, showing desired HDR with donor templates.

## Notes

### Competing Interest Statement

The authors have declared no competing interest.

